# Dynamic Epigenetic Changes During Antidepressant Pharmacotherapy in Major Depressive Disorder

**DOI:** 10.1101/2025.11.20.688229

**Authors:** Svenja Müller, Josef Frank, Eric Zillich, Fabian Streit, Valentin Nicolai Koch, Miriam A. Schiele, Katharina Domschke, Andre Tadic, David P. Herzog, Stefanie Wagner, Marianne B. Müller, Giulia Treccani, Nils C. Gassen, Marcella Rietschel, Stephanie H. Witt, Klaus Lieb, Jan Engelmann, Lea Zillich

**Affiliations:** Department of Genetic Epidemiology in Psychiatry, Central Institute of Mental Health, Medical Faculty Mannheim, Heidelberg University, Mannheim, Germany; German Center for Mental Health (DZPG), partner site Mannheim/Heidelberg/Ulm, Germany; Department of Ergonomics, Leibniz Research Centre for Working Environment and Human Factors (IfADo), Dortmund, Germany; Hector Institute for Artificial Intelligence in Psychiatry (HITKIP), Central Institute of Mental Health, Medical Faculty Mannheim, Heidelberg University, Mannheim, Germany; Department of Psychiatry and Psychotherapy, Medical Center-University of Freiburg, Faculty of Medicine, University of Freiburg, Freiburg, Germany; German Center for Mental Health (DZPG), Partner Site Berlin/Potsdam, Berlin, Germany; Department of Psychiatry and Psychotherapy, University Medical Center, Mainz, Germany; Department of Psychiatry, Psychotherapy and Psychosomatics, Dr. Fontheim Mentale Gesundheit, Liebenburg, Germany; Translational Psychiatry, Department of Psychiatry and Psychotherapy & Focus Program Translational Neuroscience, University Medical Center, Mainz, Germany; Department of Systemic Neuroscience Institute of Anatomy and Cell Biology, Philipps University Marburg, Marburg, Germany; Neurohomeostasis Research Group, Department of Psychiatry and Psychotherapy, University Hospital Bonn, Bonn, Germany; Center for Innovative Psychiatric and Psychotherapeutic Research, Biobank, Central Institute of Mental Health, Medical Faculty Mannheim, Heidelberg University, Mannheim, Germany

**Author notes:** Correspondence: Dr. Lea Zillich; Department of Psychiatry and Psychotherapy; University Medical Center Freiburg; Hauptstr. 5; 79104 Freiburg, Dr. Jan Engelmann; Department of Psychiatry and Psychotherapy; University Medical Center Mainz; Untere Zahlbacher Str. 8; 55131 Mainz. Shared senior authorship: Jan Engelmann, Lea Zillich.

## Abstract

Although antidepressants remain the main pharmacological treatment for major depressive disorder (MDD), therapeutic response varies, highlighting the need for molecular markers that predict treatment response, as well as insights into the biological processes underlying antidepressant efficacy. Candidate-gene and cross-sectional epigenome-wide association studies (EWAS) have reported DNA methylation signatures associated with antidepressant response; however, findings remain inconsistent. To date, no longitudinal EWAS has examined methylation trajectories across multiple time points during antidepressant treatment in MDD. Within the Early Medication Change trial, DNA methylation data was generated for 162 patients with MDD (81 responders, 81 non-responders) and 48 matched healthy controls. Patients were assessed at four times across eight-weeks of standardized antidepressant treatment, while controls were assessed twice. Differentially methylated positions (DMPs) and regions (DMRs) were identified using longitudinal EWAS models and the comb-p algorithm. Baseline methylation levels were associated with depressive symptom severity at day 28 and day 56 through two and six DMRs, respectively (e.g., *TNRC6C*, *CAT*), whereas no single DMP reached significance. Longitudinal analyses identified one DMP associated with improvement in depressive symptoms (*YLMP1*) and three DMRs showing methylation changes over time (e.g., *GPR126, PM20D1)*. Combined patient-control analyses revealed additional DMPs and DMRs associated with diagnosis and temporal effects. This study provides first longitudinal evidence of regionally coordinated DNA methylation changes during antidepressant pharmacotherapy, revealing alterations in genes involved in neuroplasticity and inflammatory processes that are associated with clinical response. Replication and functional validation will be essential to determine their relevance for personalized antidepressant treatment.

## Introduction

Major depressive disorder (MDD) affects over 300 million people globally and is a leading cause of disability worldwide (1–3). Pharmacological treatment with antidepressants is recommended for moderate to severe depressive episodes (4–6). However, treatment response is highly variable, often necessitating medication adjustments (7,8). Investigating biological changes associated with antidepressant response may help to understand the underlying mechanisms of both depression and treatment efficacy.

MDD is increasingly understood as a disorder of dysregulated neuroplasticity rather than solely of neurotransmitter imbalance (9,10). Research has emphasized the critical role of intracellular signaling, gene expression, and protein translation as key mediators of antidepressant action (10). Antidepressants promote synaptic connectivity within and between neuronal networks that regulate depression-related behaviours (11). Within this framework, epigenetic modifications, particularly DNA methylation, has emerged as a dynamic process through genetic predisposition and environmental factors (e.g. medication) modulate neuroplasticity, stress responsiveness and emotional regulation (12–14). Importantly, DNA methylation can be highly dynamic, with changes occurring within minutes to hours, thereby contributing to rapid alterations in gene and protein expression that support short-term neuronal adaptations and experience-dependent plasticity (15). This underlines the potential role of DNA methylation as a molecular signature of antidepressant response. Although DNA methylation is highly tissue-specific, peripheral DNA methylation can serve as a correlate for brain-related processes (15,16) and is ideally suited for clinical translation.

Previous epigenetic investigations of antidepressant treatment response mainly focused on selected target genes like *BDNF, 5-HTT* and *FKBP5 (e.g.* 17–21), or the serotonergic and noradrenergic pathways (reviews: 22,23). Beyond candidate approaches, our group and others have conducted pre-treatment, single-time-point epigenome-wide association studies (EWAS) to explore epigenetic signatures associated with pharmacological antidepressant treatment response. Among others, Ju et al. identified several CpG sites differentially methylated between responders (N = 82) and non-responders (N = 95) after eight weeks of treatment, including a DMP within the *CHN2* gene that showed the strongest association with mRNA expression (24). In our EWAS comparing responders and non-responders (N = 40 each), no epigenome-wide significant DMP was detected, but 20 DMRs were found near genes such as *SORBS2* and *CYP2C18* (*25*). Recently, Schiele et al. (26) reported suggestive associations emerged at CpG sites linked to genes including *TIMP2, VDAC1, and SORL1* after six weeks of antidepressant treatment (N = 230). Despite the dynamic nature of DNA methylation patterns and evidence that antidepressants may exert their effects by modifying these epigenetic patterns (27,28), longitudinal studies are rare and have largely focused on candidate genes (29,30). Regarding psychotherapeutic treatment, recent research on trauma-focused therapies, including cognitive-behavioral therapy or Eye Movement Desensitization and Reprocessing, reported 110 DMRs following treatment in individuals with treatment-resistant depression (31). So far, a limited number of studies have investigated other non-pharmacological interventions. For instance, Sirignano et al (32) identified a single differentially methylated site within *TNKS* associated with response to electroconvulsive therapy (ECT). However, to the best of our knowledge, there are currently no studies that have systematically and longitudinally investigated epigenome-wide methylation changes throughout a pharmacological antidepressant treatment, from treatment initiation through response to clinical remission.

To address this gap, we performed an epigenome-wide analysis of DNA methylation alterations associated with antidepressant treatment response. We compared epigenetic profiles among responders, age- and sex-matched non-responders, and matched healthy controls by assessing DNA methylation differences prior to treatment initiation and at three additional time points throughout an eight-week standardized antidepressant treatment. All participants were recruited from the large, well-characterized Early Medication Change (EMC) cohort (33,34). This longitudinal design allowed us to examine two complementary aspects: first, the predictive value of baseline DNA methylation profiles for treatment outcome; and second, dynamic changes in DNA methylation trajectories during treatment, providing mechanistic insights into the molecular processes underlying antidepressant response. By capturing both baseline epigenetic states and their trajectories, we aimed to identify DNA methylation patterns associated with treatment outcome and potential molecular targets. Comparing patients to healthy controls (assessed twice) allowed characterization of alterations beyond medication effects. Together, this study provides comprehensive insights into molecular signatures associated with antidepressant response, informing the development of biomarkers and precision medicine approaches for the treatment of MDD.

## Materials and Methods

### Study cohort

This investigation is a secondary analysis of 162 MDD patients from the ’Randomised Clinical Trial Comparing an Early Medication Change (EMC) Strategy with Treatment as Usual (TAU) in MDD’ (ClinicalTrials.gov, NCT00974155). Out of 889 patients originally enrolled (between 2009 and 2014), genetic data was available for 560 (35). Of these, 81 responders and matched non-responders were selected based on their treatment response at day 28. Responders showed at least a 50% improvement in depressive symptoms, as measured by the 17-item Hamilton Rating Scale for Depression (HAMD-17), after four weeks of treatment according to study protocol. In addition, 48 matched healthy controls were available and included in this investigation (36). The detailed study design is shown in Fig. 1.

**Figure 1.**
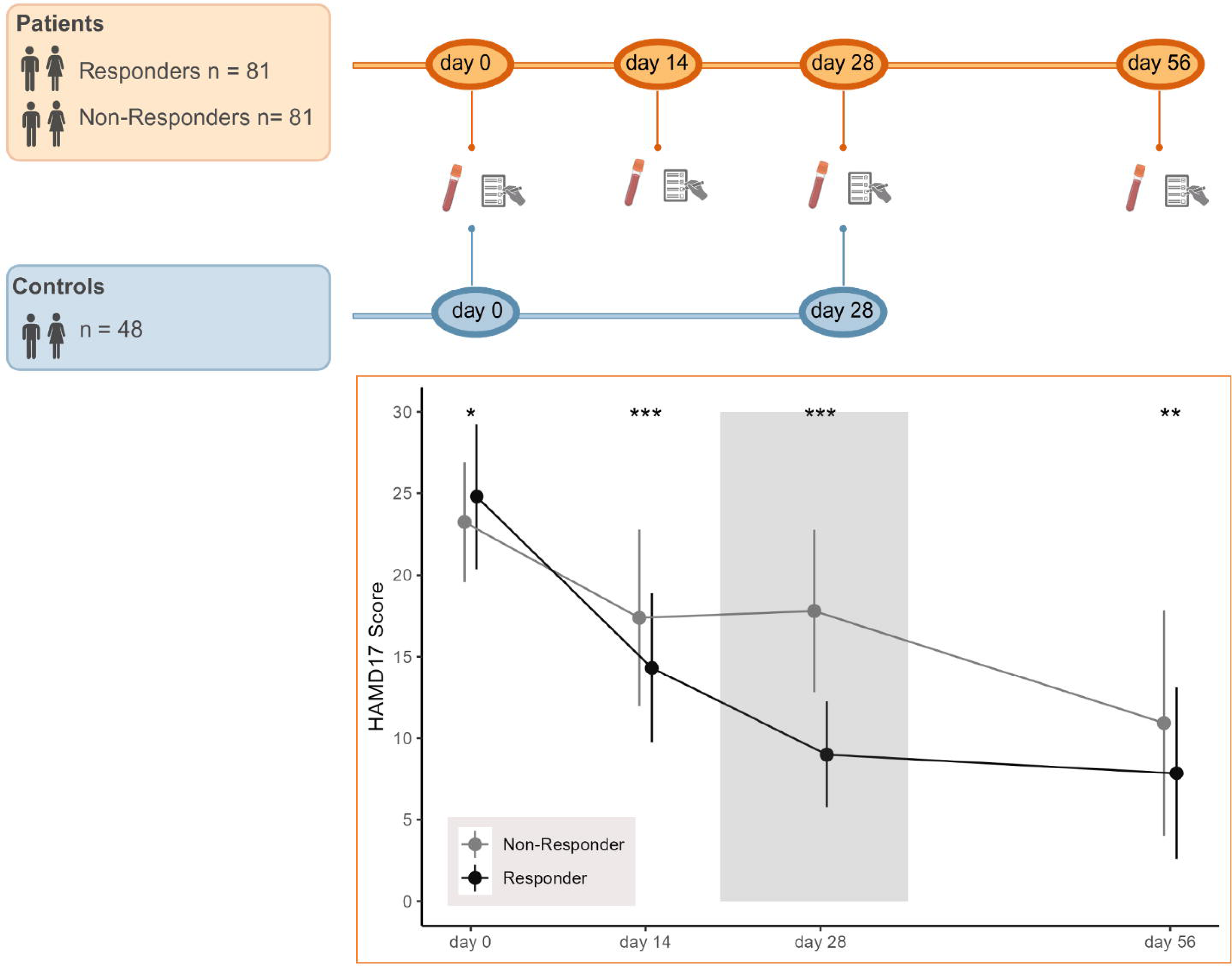
Study design and Time Course of HAMD17 Scores in Responders and Non-Responders. A. Overview of blood sampling and HAMD-17 assessment time points in patients and controls. B. Time course of depressive symptom severity measured by the Hamilton Depression Rating Scale (HAMD17) in responders and non-responders over 56 days. Means and standard deviation are given. HAMD scores were significantly different between groups at all time points (p < .05). At Day 28, responders showed a ≥50% reduction in symptom severity compared to baseline, whereas non-responders displayed only a modest decline in symptom scores (<50%). Statistical significance is indicated as follows: *p* < 0.05 (\**), p < 0.01 (**), and p < 0.001 (\*\*\**).

The EMC trial, conducted between 2009 and 2014, was a multicenter, randomized controlled clinical trial designed to evaluate whether patients who do not show improvement after 14 days of antidepressant treatment with escitalopram benefit from an early medication change. In this protocol, non-responders at day 14 were transitioned to venlafaxine. Those who continued to show no response at day 28 received an augmentation with lithium. This accelerated treatment strategy was compared against patients treated according to current guideline recommendations (treatment as usual; TAU), in which escitalopram was continued for an additional two weeks, and patients were only switched to venlafaxine in the case of non-response from day 29 onwards. All participants provided written informed consent after a complete and extensive description. The study was approved by the ethics committee of the Landesärztekammer Rheinland-Pfalz and was conducted in accordance with the Declaration of Helsinki. Details of the study protocol and the inclusion/exclusion criteria have been reported elsewhere (33,34). The controls were recruited as part of a subproject of the EMC trial through posters in the study center.

### Study procedures

Diagnoses were verified at screening using the German versions of the Mini International Neuropsychiatric Interview (M.I.N.I.) and the Structured Clinical Interview for DSM-IV Axis II Personality Disorders (SCID-II). Sociodemographic and clinical data were obtained via patients‘ self-report. Depression severity was assessed weekly from baseline to day 56 using the HAMD17 by trained and blinded raters. If necessary, antidepressant premedication was washed out following inclusion and prior to the baseline visit. According to the study protocol, all participants received escitalopram (20 mg/day) from baseline to day 14, followed by a predefined treatment algorithm. The used antidepressant treatment algorithm can be openly accessed at https://trialsjournal.biomedcentral.com/articles/10.1186/1745-6215-11-21. Other antidepressants, antipsychotics, mood stabilizers, or medications with a substantial impact on the primary study endpoints were not permitted. Concomitant pharmacotherapy was restricted to the treatment of transient symptoms or medication-related side effects and included short-acting hypnotics (zolpidem or zopiclone), low-potency antipsychotics (pipamperone), promethazine at standard doses, and benzodiazepines up to a diazepam-equivalent dose of 15 mg. Treatment for general medical conditions was permitted and remained unchanged provided that these medications did not contraindicate the use of the study medication (33,34).

### Quantification of DNA methylation

Genomic DNA was extracted from whole blood using the QIAamp DNA Blood Midi Kit (Qiagen, Hilden, Germany). Blood samples were collected in fasting patients in the morning before medication intake at baseline (prior to start of study medication) and at day 14, 28 and 56 (see Fig. 1). For control participants, samples were obtained at baseline and at day 28. All extracted DNA samples were stored at −20 °C until further processing. Responders and non-responders were matched by age and sex, and DNA from these matched pairs, along with samples from controls, was randomized and pipetted onto processing plates. A total of 500 ng of genomic DNA per sample was subjected to bisulfite conversion using the EZ-96 DNA Methylation-Gold Kit (Zymo Research, Irvine, USA). Genome-wide DNA methylation profiling was performed using the Infinium HumanMethylationEPIC BeadChip array, with scanning conducted on the Illumina HiScan system (Illumina, San Diego, CA) at the Genome Analysis Center at the Helmholtz Institute München.

### Data preprocessing, quality control, and filtering

All analyses were conducted using R version 4.4.1. (37). Methylation data were extracted from raw signal intensity files using an updated version of the CPACOR pipeline (38,39). Standard sample- and probe-level quality control procedures were applied, including the exclusion of samples failing quality control (QC) and probes with low reliability. After preprocessing, 164 patients (with baseline and at least one follow-up time point) and 51 control participants (43 of whom had data at both time points) were retained for analysis. A total of 642 148 CpG sites passed QC and remained for downstream analyses. Detailed preprocessing steps and QC thresholds are provided in Supplementary Information S1.

To account for technical variability and batch effects, principal component analysis (PCA) was performed on the internal control probes of the EPIC array. The first four principal components were included as covariates in all subsequent analyses. Cell-type heterogeneity was estimated using a reference-based deconvolution approach based on the method of (40), which provides six proportional estimates summing approximately to one. To avoid multicollinearity in the EWAS, variance inflation factors were calculated for each estimate, and the granulocyte proportion - identified as contributing to collinearity - was excluded from further models.

### Covariates

All analyses were adjusted for relevant covariates, including sex, age, smoking status (approximated via DNA methylation at *AHRR* cg05575921; 41), and body mass index (BMI). For two individuals with missing BMI values, group- and sex-specific mean imputation was applied. To control for cellular heterogeneity, estimated proportions of major blood cell types (CD8+ T cells, CD4+ T cells, monocytes, B cells, and NK cells) were included. In addition, two genotype-based principal components (PCs) were added to account for population stratification, and four control probe PCs were included to correct for batch effects. Samples lacking genotype information or excluded during genotype QC (n=2 patients, n=3 controls) were removed from analyses, yielding a final sample of 162 patients and 48 controls. For details on genotype QC, see Supplementary information S2. To evaluate the relative contribution of each covariate to the variance in DNA methylation and to justify their inclusion in downstream models, we performed variance partitioning using linear mixed models (implemented in the *variancePartition* R package; 42).

### Epigenome-wide association study

The final dataset, comprising 162 patients and 48 controls after preprocessing and QC, was analyzed using linear models (LM) for the cross-sectional methylation analyses and linear mixed models (LMMs) implemented in the *lme4* R package (43) for the longitudinal methylation analyses. Three separate models were used to address distinct research questions. All models accounted for symptom severity at baseline. Model 1 achieved this by including baseline HAMD, whereas Models 2 and 3 incorporated HAMD as a time-varying predictor. Model 1 used linear regression to predict HAMD scores at the final time point (day 56) based on methylation levels and HAMD scores at baseline. In parallel, we applied the same approach to HAMD scores at the clinically relevant mid-treatment assessment (day 28), which forms the basis for our responder vs. non-responder classification. This approach (including 642 148 sites) allowed us to assess whether early methylation patterns and symptom severity could predict later clinical outcomes. Multiple testing correction was applied using the False Discovery Rate (FDR) approach according to Benjamini and Hochberg (44) and the resulting values are reported as q-values. CpG sites were annotated according to the manufacturer’s manifest (Illumina MethylationEPIC v1 B5). Model 2 focused on patients (n=162) and included four time points. The model assessed the association between DNA methylation and symptom severity, while accounting for within-subject correlation across time via random intercepts. For each CpG site, the effects of time (day 14, day 28, and day 56 relative to baseline) and HAMD score (continuous) were estimated. Estimated marginal means (EMMs) were computed using the *emmeans* R package (45) to derive average methylation levels at each time point. After model singularity checks, 606 546 CpG sites remained for this analysis. Model 3 examined longitudinal methylation changes from baseline to day 28 in patients (n = 162) and controls (n = 48). For each CpG site, the effects of time, HAMD score (continuous), and diagnostic group (patients vs. controls) were estimated. After model singularity checks, 562 713 CpG sites remained for this analysis. To assess bias and inflation in test statistics across all models, we calculated quantile-quantile (QQ) plots and estimated the inflation factor lambda (λ). Bacon correction using the Bayesian method implemented in the *bacon* R package (46) was applied for model 3 comparing patients and controls, where λ was elevated (λ = 1.55). Full methodological details of QC procedures and bacon correction are provided in Supplementary Information S3.

### Downstream analyses

Differentially methylated regions (DMRs) were identified using the comb-p function from the *ENmix* R package (47). This approach accounts for spatial autocorrelation among neighboring CpG sites by aggregating adjacent significant sites into regions of methylation enrichment (48). A sliding window was applied with a maximum inter-CpG distance of 750 base pairs and a bin size of 500 base pairs. Regions were initiated using a seed p-value threshold of 0.001. Multiple testing correction across identified regions was performed using the Sidak method. Only DMRs containing at least two CpG sites were retained for downstream analysis.

Gene set enrichment analysis was conducted using the gometh function from the *missMethyl* R package to identify overrepresented Gene Ontology (GO) biological processes among differentially methylated CpG sites (49). The analysis was based on CpG sites with a nominal p-value < 0.001. The full set of tested CpG sites was used as the background, and the Illumina EPIC array annotation was specified.

## Results

### Sample characteristics

The clinical sample comprised 49.4% women, with a mean age of 41.7 years (SD = 11.4). Treatment responders were more likely to present with a first depressive episode, whereas non-responders had a higher prevalence of recurrent depression (p = 0.013). Detailed sociodemographic and clinical characteristics are summarized in Table 1. In accordance with the study protocol, all patients were treated with escitalopram at a fixed dose of 20 mg/day during the initial two-weeks. Patients classified as responders in this analysis, defined by continuous improvement in depressive symptomatology over time, were subsequently maintained on escitalopram according to the EMC randomization scheme, with majority (88.9%) continuing 20mg escitalopram till end of observation. In contrast, non-responders were treated according to the predefined treatment algorithm and were predominantly switched to high-dose venlafaxine after week 4 due to non-response (82.7%). Responder and non-responder groups differed significantly in symptom severity at all time points, with the smallest difference at baseline and progressively larger differences at day 14, day 28, and day 56 (see Figure 1). The control group included 60.4 % women, with a mean age of 38.3 years (SD = 12.8), a mean BMI of 23.7 (SD = 3.7), and 45.8 % reporting current smoking. Among sociodemographic variables, BMI was the only significant difference between patients and controls, being higher in patients (p < 0.001).

**Table 1.** Clinical and sociodemographic data.

|  | Total<br>(n = 162) | Responders<br>(n = 81) | Non-Responders<br>(n = 81) | p-value (group<br>comparison) |
| --- | --- | --- | --- | --- |
| Age - yrs (SD) | 41.71 (11.43) | 41.79 (11.28) | 41.63 (11.66) | 0.929<sup>1</sup> |
| Sex female (%) | 80 (49.4 %) | 41 (50.6 %) | 39 (48.1 %) | 0.875<sup>2</sup> |
| BMI (SD) | 27.04 (6.5) | 26.47 (6.25) | 27.62 (6.25) | 0.262<sup>1</sup> |
| Smoking – yes (%) | 50 (31.3 %) | 30 (37.0 %) | 20 (25.3 %) | 0.153<sup>2</sup> |
| Age at onset – yrs (SD) | 33.65 (12.58) | 34.9 (12.37) | 32.4 (12.7) | 0.206<sup>1</sup> |
| Duration of current episode<br>– weeks (SD) | 32.29 (39.69) | 34.15 (38.63) | 30.43 (40.88) | 0.553<sup>1</sup> |
| Course of disorder (%) |  |  |  | **0.013<sup>2</sup>** |
| First episode | 56 (34.6 %) | 36 (44.4 %) | 20 (24.7 %) |  |
| Recurrent | 106 (65.4 %) | 45 (55.6 %) | 61 (75.3 %) |  |

*Note:* Continuous measures are presented as mean (standard deviation) and categorical measures as frequency (percent). M mean, SD standard deviation, yrs years. Response was defined as a decrease in HAMD17 from baseline to day 28 of $\geq 50\%$ ; 1 = *t*-Test (or Welch test in case of variance inequality); 2 = $\chi^2$ test.

### Baseline methylation associated with treatment outcome

In the models predicting depressive symptom severity at day 28 and day 56, respectively, no epigenome-wide DMP reached significance after FDR-correction for multiple testing. The regression coefficients corresponding to the 100 DMPs most strongly associated with treatment outcomes are provided in Supplementary Tables S1-S2. Using regional analysis, we identified six DMRs whose methylation levels were associated with symptom severity at day 56 and two DMRs associated with symptom severity at day 28. The DMR showing the strongest (positive) association with symptom severity at day 56 was annotated to *Trinucleotide Repeat Containing 6C* (*TNRC6C*), a protein-coding gene; a largely overlapping DMR was also observed at day 28. Figure 2 shows the Manhattan plot highlighting DMRs associated with symptom severity at day 56 in green, and the corresponding plot for the daylJ28 analysis is provided in Supplementary Figure S1. Details on the DMRs identified in both analyses are provided in Supplementary Table S3.

**Figure 2.**
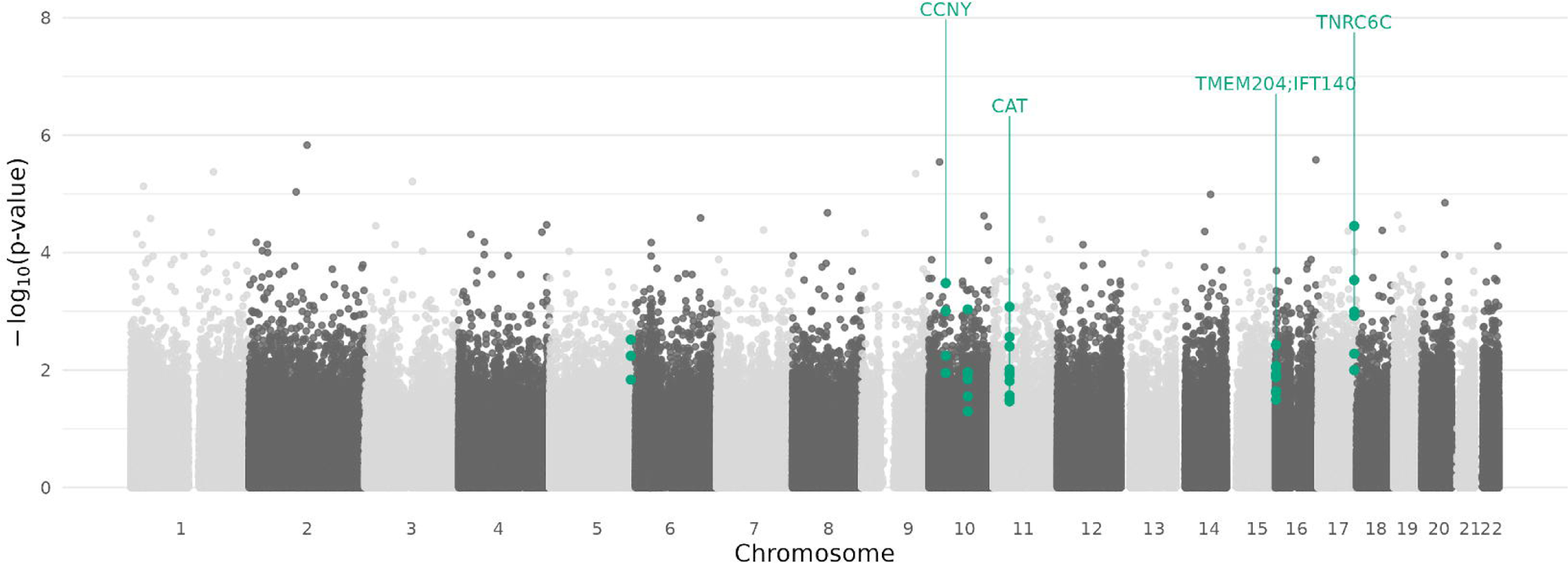
Manhattan plot illustrating the association between DNA methylation and depressive symptom severity at day 56. Shown are –log_₁₀_(p-values) across the genome for all tested CpG sites. Differentially methylated regions identified in the regional analysis are highlighted in green.

### Longitudinal DNA methylation changes and associations with symptom improvement in patients

The linear mixed-effects model examined the effects of time (day 14, day 28, and day 56 relative to baseline) and continuous symptom change across the eight-week treatment period. Although the original sample was selected based on matched responder and non-responder groups, the model assessed longitudinal changes in HAMD scores across all patients, thereby leveraging the repeated-measures design. While no DMP reached significance for the effect of time, one epigenome-wide DMP was significant after FDR-correction for the effect of symptom change. Figure 3 illustrates associations between DNA methylation levels and symptom improvement across the treatment period, while Supplementary Figures S2–S4 show the time-specific effects for day 14, day 28, and day 56 compared to baseline. A single significant DMP (cg15731816) associated with HAMD score was annotated to *YLP Motif Containing 1* (*YLPM1;* β = 0.0005, p = 1.25 x 10^-8^, q = 0.008), with the positive beta indicating that increased methylation levels corresponded to higher HAMD scores over time. Importantly, over the course of treatment, both methylation at this site and HAMD scores generally decreased, suggesting that reductions in methylation were linked to clinical improvement (see Fig. 4). Supplementary Tables S4-S7 present the 100 DMPs most strongly associated with continuous changes in depressive symptoms across treatment and with time-specific effects, respectively. Regional analyses revealed three DMRs in which methylation levels changed significantly over the course of treatment from baseline to day 28, and one DMR from baseline to day 56. Of note, the same DMR – including five CpG sites – was detected at both day 28 and day 56 and mapped to *G protein-coupled receptor 126* (*GPR126)*, a protein-coding gene. Details on the DMRs are provided in Supplementary Table S8.

**Figure 3.**
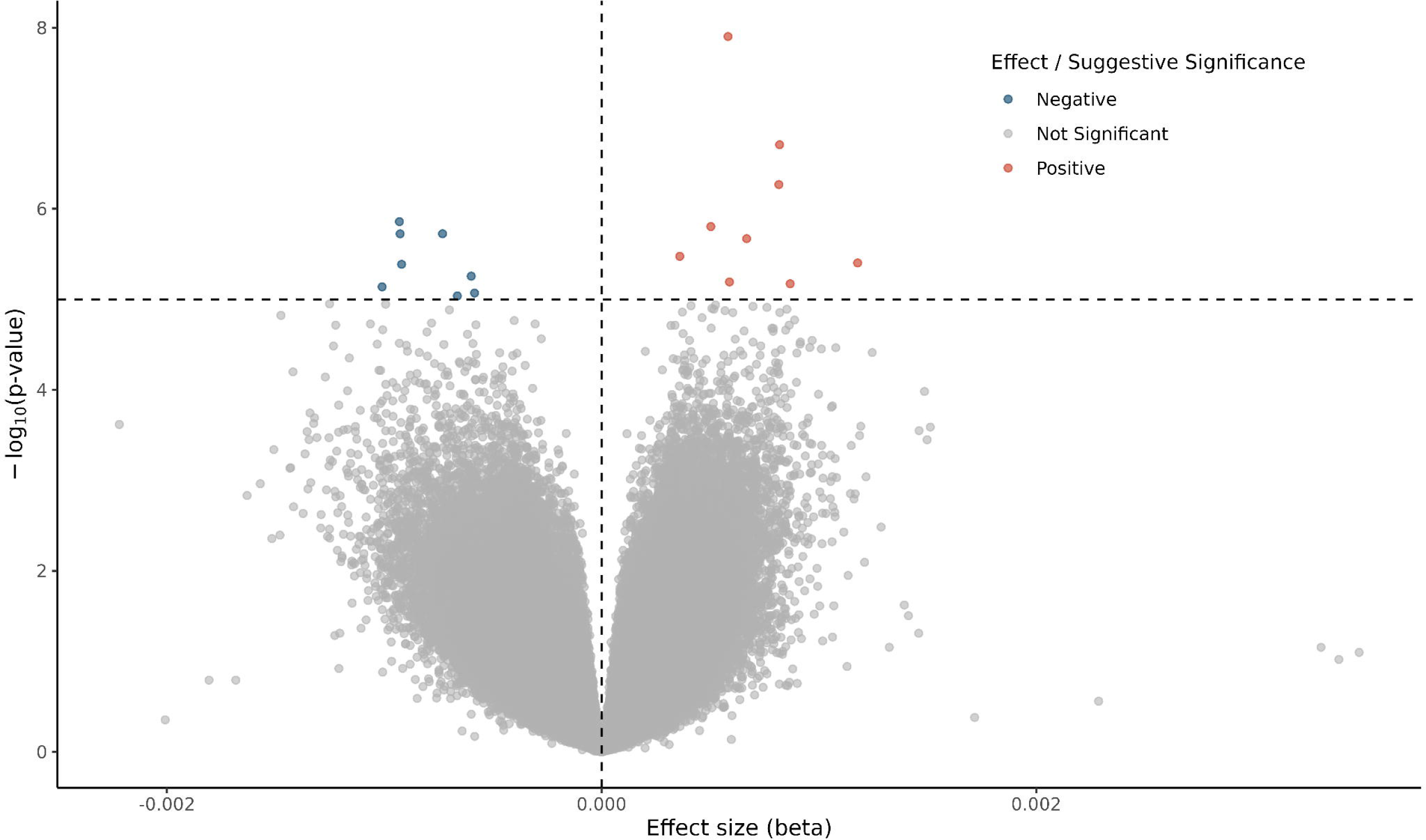
Volcano plot illustrating the associations between DNA methylation levels and continuous changes in depressive symptom severity in patients. Each point represents a CpG site, with the x-axis showing the effect size (*β*) and the y-axis the –log_₁₀_-transformed p-value. CpG sites with a positive association (increased DNA methylation with higher symptom severity) are shown in red, and negative associations are shown in blue. Horizontal and vertical dashed lines indicate the thresholds for suggestive statistical significance (p < 1 × 10^-4^) and effect size, respectively.

**Figure 4.**
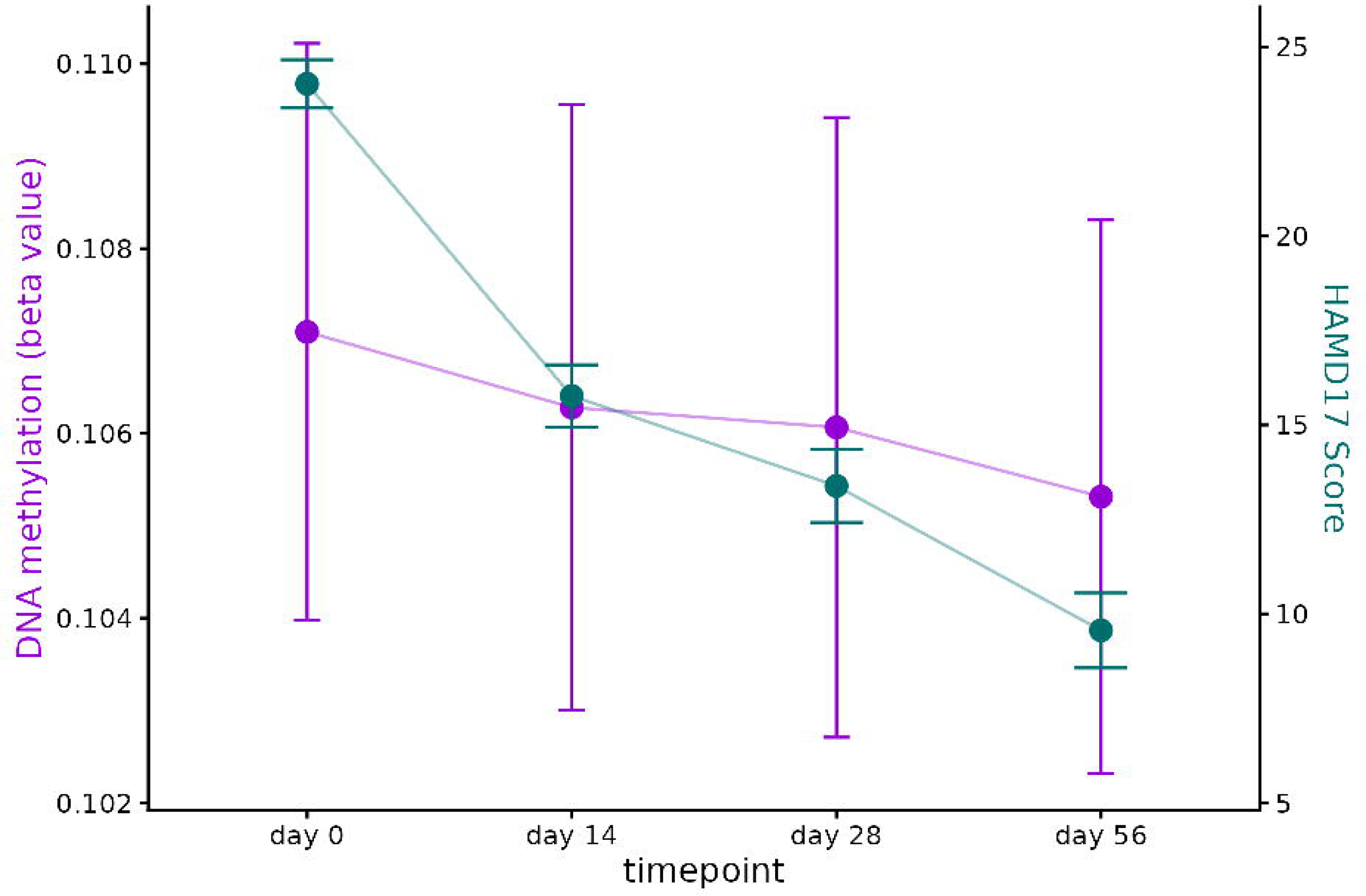
Longitudinal trajectories of HAMD scores and DNA methylation at cg15731816 (in patients). Mean Hamilton Depression Rating Scale (HAMD17) scores (right y-axis) and DNA methylation at cg15731816 expressed as *β* values (left y-axis) across four time points (n=162). Points represent means; error bars indicate 95% confidence intervals.

### Epigenetic differences between patients and controls across two time points

In the linear-mixed model comparing patients and controls across two time-points, four CpG sites were identified as significantly differentially methylated for the main effect of group (patients vs. controls; see Table 2). The most pronounced difference was observed at cg17232014 in *Heme-binding protein 1* (*HEBP1;* β *=* 0.04, p= 2.6 x 10^-8^, q = 0.015). No CpG sites showed statistically significant methylation differences for the effect of time (day 28 relative to baseline) or symptom change (change in HAMD score from baseline to day 28) after multiple testing correction. The regression coefficients corresponding to the 100 DMPs most strongly associated with group, time point, and symptom change are provided in Supplementary Table S9-S11, respectively. The subsequent DMR analysis identified four DMRs associated with time point, one DMRs associated with HAMD score, but none with group. The DMR showing the strongest association with time was annotated to the protein-coding gene *Tenascin XB* (*TNXB)* and showed an increase in methylation from baseline to day 28. See Table 2 for a list of the identified DMPs and DMRs.

**Table 2.** Differentially methylated sites and regions associated with time (day 28 relative to baseline), symptom severity, and group differences (patients vs. controls)

| **A.** |  |  |  |  |  |  |  |  |
| --- | --- | --- | --- | --- | --- | --- | --- | --- |
| **effect (DMPs)** | **chr** | **position** | **p_corr** | **fdr p** | **probe id** | **Gene** | **beta** |  |
| group | 12 | 13153193 | 2.60E-08 | 0.014 | cg17232014 | *HEBP1;*<br>*HTR7P1* | 0,0396 |  |
|  | 2 | 3142407 | 1.50E-07 | 0.028 | cg08902894 |  | -0,0249 |  |
|  | 18 | 77106426 | 1.40E-07 | 0.028 | cg27410534 | *ATP9B* | -0,0226 |  |
|  | 1 | 32739049 | 3.35E-07 | 0.047 | cg10325497 | *LCK* | -0,0201 |  |
| **B.** |  |  |  |  |  |  |  |  |
| **effect (DMRs)** | **chr** | **start** | **end** | **p** | **sidak p** | **n probe** | **Gene** | **Direction** |
| HAMD | 6 | 32847441 | 32847831 | 3.39E-09 | 4.89E-06 | 11 | *PPP1R2P1* | + |
| day 28 | 6 | 32063403 | 32064162 | 1.81E-10 | 1.34E-07 | 19 | *TNXB* | + |
|  | 15 | 66947167 | 66947618 | 2.31E-08 | 2.88E-05 | 6 | *LINC01169* | + |
|  | 6 | 32847441 | 32847831 | 1.32E-08 | 1.91E-05 | 11 | *PPP1R2P1* | + |
|  | 5 | 8457599 | 8458090 | 1.99E-07 | 0.0002 | 6 | *MIR4458HG;*<br>*LOC729506* | + |

Note: chr Chromosome, + indicates higher methylation is associated with higher HAMD scores (upper part: HAMD), + indicates increasing methylation over the specified time period (lower part: day 28)

### Gene-set enrichment analyses

In the gene-set enrichment analyses, several gene ontology categories showed nominally significant p-values across the different models. For example, in the second model (longitudinal EWAS in patients only), the biological process “regulation of microvillus assembly” associated with time (day 56 relative to baseline; p = 6.4 × 10^-4^, q = 1) exhibited a strong nominal signal, alongside specific molecular functions related to transcription and protein modification (i.e., “RNA polymerase II core promoter sequence-specific DNA binding” associated with time [day 28 relative to baseline; p = 4.61 × 10^-5^, q = 1]; “protein methyltransferase activity” associated with continuous change in depressive symptom severity [p = 3.7 × 10^-4^, q = 1]). However, none of these findings remained statistically significant after correction for multiple testing. Results of the gene-set enrichment analyses are provided in Supplementary Tables S12-S20.

## Discussion

Previous epigenetic studies in MDD have primarily examined baseline DNA methylation levels as a potential predictor of antidepressant response, yielding mixed findings and providing limited insight into within-subject dynamics during treatment. Most prior work was cross-sectional or restricted to candidate genes, thus failing to capture the temporal trajectory of methylation patterns during pharmacotherapy. Longitudinal methylome-wide investigations have mainly focused on non-pharmacological interventions, such as ECT or psychotherapy, leaving the epigenetic dynamics of standardized antidepressant treatment largely unexplored. To address this gap, the present study profiled genome-wide methylation at baseline and at three follow-up time points over an eight-week course of antidepressant treatment, comparing responders, non-responders, and non-depressed healthy controls within a randomized treatment framework.

Although the current EWAS did not identify any individual DMPs at baseline significantly associated with HAMD score at day 28 or day 56, respectively, six DMRs were positively related to depressive symptom severity at day 56. The top DMR was annotated to *TNRC6C*, a key component of microRNA-mediated gene silencing, suggesting potential involvement of post-transcriptional regulatory mechanisms in antidepressant response, and previously implicated in psychiatric (50) and cardiovascular disease (51). Prior EWAS have identified single-CpG associations with antidepressant response in genes such as *CHN2* (24). In our previous analysis, we identified 20 DMRs, notably in *SORBS2* and *CYP2C18*. Similarly, Schiele et al. (26) found no epigenome-wide significant CpGs predicting response after six weeks of treatment but reported suggestive associations in genes such as *TIMP2*, *VDAC1*, and *SORL1*, which have been implicated in mitochondrial function and extracellular matrix regulation (52,53). Notably, variability across pre-treatment EWAS likely reflects differences in sample size, analytical pipelines, cohort characteristics, and medication regimens. Future meta-analytic efforts integrating longitudinal EWAS and transcriptomic data, combined with cell-type-specific and functional annotations, will be crucial to enhance reproducibility and clarify the mechanistic underpinnings of antidepressant response.

While identifying pre-treatment DNA methylation profiles for antidepressant response is crucial from a predictive perspective, longitudinal assessments offer complementary insights into methylation trajectories associated with treatment outcomes and may reveal potential mechanistic targets. Using a linear mixed model that accounted for repeated measures within individuals, we identified one CpG site (cg15731816, annotated to *YLPM1*) that was significantly associated with changes in depressive symptom severity across the treatment period. Specifically, higher methylation levels at this site were linked to higher symptom severity, while both methylation levels and HAMD scores generally decreased over time, suggesting that reductions in methylation levels may reflect or contribute to clinical improvement. Interestingly, a recent pan-ancestry analysis of sequencing data from three large biobanks identified core genes underlying a broad spectrum of human diseases, including *YLPM1*, which was implicated in psychiatric disease (54). Complementary evidence from cell type-specific expression quantitative trait locus (eQTL) analyses in MDD highlighted *YLPM1* as a candidate gene in oligodendrocytes, emphasizing the role of cellular heterogeneity in MDD pathogenesis (55). Moreover, a cross-ancestry rare variant study of obesity identified *YLPM1* as a gene expressed in both brain and adipose tissue, associated with obesity traits and neuropsychiatric phenotypes, pointing to potential pleiotropic effects on metabolism and mental health (56). Taken together, these converging lines of evidence support a role for *YLPM1* in modulating symptom trajectories in depression. In our study, methylation was measured in blood, but the correspondence with cell type-specific brain expression suggests peripheral methylation changes may reflect or serve as a proxy for processes relevant to symptom change. These findings highlight *YLPM1* as a potential target for stratified treatment approaches, although mechanistic studies are needed to establish causality.

The regional analysis identified dynamic DMRs, showing significant methylation changes relative to baseline at day 28 (three DMRs) and day 56 (one DMR), highlighting time-dependent effects during antidepressant treatment. One DMR annotated to *GPR126* was obtained across both time points, suggesting continuous, time-dependent epigenetic modifications throughout treatment. *GPR126*, also known as *Adhesion G protein-coupled receptor G6* (*ADGRG6),* is expressed in multiple tissues and regulates neural development via G-protein- and N-terminus-dependent signaling (57,58). The observed methylation changes in *GPR126* across the course of treatment imply ongoing modulation of adhesion G protein-coupled receptor signaling, which may indirectly reflect neural development and synaptic remodeling, supporting the conceptualization of MDD as a disorder of dysregulated neuroplasticity (9,10). Another DMR in *Peptidase M20 Domain Containing 1* (*PM20D1)* was detected at day 28, further confirming time-related methylation effects in MDD patients.

*PM20D1* has recently been highlighted in a cross-trait meta-analysis as functionally implicated in both MDD and type 2 diabetes (59). Genes associated with both mood and metabolic disorders were also reported in our previous study (e.g., *SORBS2*; 25). Altered methylation in these genes may contribute to comorbidity, potentially via inflammatory mechanisms (60). Together, these results highlight the value of longitudinal assessment and, to our knowledge, provide the first evidence of dynamic DNA methylation trajectories during antidepressant therapy. Although no methylation changes were observed at day 14, early modifications cannot be excluded, consistent with evidence that epigenetic changes are highly dynamic and can rapidly influence synaptic plasticity (61). Changes became evident from day 28 onward, emphasizing the importance of multiple time points to capture the full trajectory of epigenetic adaptation during antidepressant treatment (62).

To further disentangle disease-related modifications from treatment effects, we compared DNA methylation profiles between patients and healthy controls over time. In the case–control analysis, four CpG sites showed significant group differences, with the strongest effect observed at cg17232014 in *HEBP1*. This robust finding indicates a persistent epigenetic distinction between patients and controls, potentially reflecting disease-related epigenetic alterations. *HEBP1* has been previously implicated in neuroinflammatory processes associated with Alzheimer’s disease (63). Notably, in a separate methylome-wide association study of self-reported antidepressant use, *HEBP1* was identified among the top ten loci (64). In a meta-analysis of two MDD case-control cohorts, Li et al. (65) identified several CpG sites annotated to *TNNT3* that were associated with MDD, along with additional DMRs annotated to genes involved in neuronal synaptic plasticity and inflammation. Although the specific sites differ from those in our study, the affected genes converge on key pathways, particularly inflammatory processes. Regional analyses identified one DMR associated with symptom severity and four regions associated with time, with the most significant region mapping to *TNXB*, a gene involved in extracellular matrix organization and potentially stress-related signaling (66). These methylation changes, observed across all participants, likely reflect general temporal effects or nonspecific biological processes rather than treatment-specific alterations. Notably, *TNXB* was previously reported as a DMR associated with antidepressant treatment response in our earlier study (25). Although the exact loci differed, these findings suggest distinct but related roles of *TNXB* in temporal methylation dynamics and treatment response, highlighting the need for further investigation.

Although several gene ontology terms showed nominally significant p-values across the three EWAS models, none survived correction for multiple testing. These results should be interpreted cautiously, as they may reflect false-positive findings, which are common in high-dimensional EWAS (67). From a biological perspective, these nominally enriched pathways align with processes implicated in neuropsychiatric disorders, including signal transduction, vascular remodeling, and epigenetic regulation. Overall, these exploratory findings highlight plausible biological systems involved in depression-related methylation dynamics but require validation in larger, independent cohorts and integration with complementary multi-omics approaches.

The EMC study provides a phenotypically well-characterized and homogeneous clinical cohort, with longitudinal DNA methylation data collected at four distinct time points during a predefined antidepressant treatment protocol and including healthy controls assessed at two time points. Nevertheless, several limitations should be noted: The sample size, although relatively robust for a longitudinal epigenetic clinical cohort, may still limit the power to detect small single-CpG effects. Although responders and non-responders already differed significantly in symptom severity at baseline, these differences were accounted for in all analyses; however, baseline imbalances may still represent a potential confounding factor. All patients underwent the same treatment schedule during the first two weeks; however, later variations in medication regimes may have influenced DNA methylation patterns, potentially confounding treatment-response-related changes. The inclusion of healthy controls is an important strength, but future studies would benefit from more balanced patient-control ratios. DNA methylation was measured in peripheral blood, which may not fully reflect brain-specific processes related to MDD. Given the longitudinal design, repeated sampling of brain tissue in living participants is not feasible, making peripheral blood the most practical and ethically acceptable approach. DNA methylation patterns are dynamic, and antidepressants may partly exert their effects by modulating these marks. Future studies should integrate DNA methylation with gene expression to investigate functional consequences, and consider additional epigenetic mechanisms, such as histone modifications (68,69).

In conclusion, this longitudinal EWAS provides novel evidence for dynamic DNA methylation changes during antidepressant treatment in patients with MDD. In addition to several significant DMRs associated with treatment outcomes in the cross-sectional analysis, longitudinal models revealed specific CpG sites and regions, most notably in *YLPM1, GPR126*, *and HEBP1*, associated with symptom severity or time. *YLPM1* emerged as the only epigenome-wide significance DMP linked to symptom trajectories over time, with reductions in methylation corresponding to clinical improvement. This finding is supported by recent evidence implicating *YLPM1* in psychiatric disease and metabolic phenotypes, suggesting pleiotropic effects on multiple biological processes relevant to depressive symptom progression. The dynamic methylation changes observed in *GPR126* are particularly noteworthy given its role in neural development and synaptic signaling, reinforcing the conceptualization of MDD as a disorder of impaired neuroplasticity. Persistent methylation differences between patients and controls, such as at *HEBP1*, further point to stable, disease-related epigenetic signatures. Together, these results highlight the value of repeated-measures, longitudinal designs to disentangle temporal, treatment-related, and disease-specific effects on the methylome. Integrating cross-sectional, longitudinal, and case–control data strengthens confidence in the results, although pathway enrichment did not survive multiple-testing correction and should be interpreted cautiously. Future research should replicate these DMP and DMR associations in larger, independent cohorts and extend these findings through harmonized meta-analyses and multi-omics integration, with the long-term goal of informing epigenetic stratification of antidepressant treatment. Combining DNA methylation with functional genomic data will be essential to clarify causal mechanisms and elucidate how epigenetic regulation interacts with gene expression and clinical trajectories to shape antidepressant response and recovery.

## Acknowledgement

The EMC trial was funded by the German Federal Ministry for Education and Research (BMBF grant no: 01 KG 0906; applicants: KL, AT); the herein presented additional investigations are not part of the funding. The BMBF had no role in the conception of the study design, in the writing of the manuscript or the decision to submit the manuscript for publication. The recruitment of the control group was funded by the German Research Foundation (“Deutsche Forschungsgemeinschaft, DFG” (funding number: WA 2970/1-1). The DFG had no role in the conception of the study design, in the writing of the manuscript or the decision to submit the manuscript for publication. Further support was provided by the German Federal Ministry of Research, Technology and Space (BMFTR) through ERANET NEURON JTC2023 (Clever; R24029FF). The funding agency had no role in the conception of the study design, in the writing of the manuscript, or in the decision to submit the manuscript for publication.

The authors are grateful to the members of the EMC Study Group, which is currently involved in the acquisition of data. The EMC Study Group consists of the following members: Univ.-Prof. Dr. Klaus Lieb, Dr. André Tadić, Univ.-Prof. Christoph Hiemke, Dr. Nadine Dreimüller, Dr. Ömür Baskaya, Dr. Danuta Krannich, Dr. Sonja Lorenz, Annette Bernius, Tillmann Weichert, Dr. Markus Lorscheider, Dr. Dipl.-Psych. Stefanie Wagner, Dipl.-Psych. Isabella Helmreich, Dipl.-Psych. Karen Grüllich, Elnaz Ostad Haji (Laboratory), Yvonne Lober (study nurse), Danuta Weichert (study nurse), cand. med. Dr. Dr. Dietrich van Calker, Dr. Sonja Gerber, Dr. Claus M. Gross, Tobias Hornig, Dipl.-Psych. Rebecca Schneibel, Ritva Atmannsbacher (Freiburg); Prof. Dr. Dieter F. Braus, Dr. Julia Reiff, Dr. Christoph Kindler, Dr. Svenja Davis, Dr. Tina Eyalil, Dr. Thorgerdur Reynisdottir, Dr. Claudia Ginap, Dipl.-Psych. Julia Kraus, Dipl.-Psych. Sabine Kaaden, Dipl.-Psych. Jelena Janzen, Dipl.-Psych. Nina Löffler, Caterina Topaloglu (Wiesbaden). The authors wish to thank Susanne Wittmann and Peter Lichtner at the Genome Analysis Center at the Helmholtz Institute München for running the methylation arrays.

## Conflict of Interest

A Tadic is designated as inventor of the European patent number 12171541.1–2404 ‘Method for predicting response or non-response to a mono-aminergic antidepressant’. He has received during the last five years consultancy fees from Janssen and ROVI. K Lieb is designated as inventor of the European patent number 12171541.1–2404 ‘Method for predicting response or non-response to a mono-aminergic antidepressant’. K Domschke is a member of the Neurotorium Editorial Board, Lundbeck Foundation, and has received speaker’s honoraria by Janssen-Cilag. All other authors declare no competing interests.

## Notes

### Summary of Updates

This version has been revised as follows: - Abstract and introduction revised for improved clarity - Additional analysis included - New figure added (Figure 4) - Tables moved to supplementary materials - Supplementary figure added - Minor corrections to statistical reporting - References updated

