## Supplementary Information, Figures for "Dynamic Epigenetic Changes During Antidepressant Pharmacotherapy in Major Depressive Disorder"

**Supplementary Information S1. DNA methylation preprocessing and quality control**

Methylation data were extracted from raw signal intensity files using an updated version of the CPACOR pipeline, including quality control procedures as described by (Lehne et al., 2015; 2016). Samples were excluded based on the following criteria: (i) poor DNA quality, defined as a missing rate >10%, or (ii) a mismatch between predicted sex (based on methylation patterns) and reported phenotypic sex. Based on these thresholds, 15 samples were excluded due to low DNA quality and an additional 20 due to sex discrepancies. Furthermore, 2 patients and 8 control participants were excluded due to missing methylation data at baseline. Probes were excluded based on the following criteria: (i) a call rate below 98%, (ii) presence of a single nucleotide polymorphism (SNP) with a minor allele frequency >10% within the probe sequence, or (iii) location on sex chromosomes (X or Y). After completion of all preprocessing and quality control steps, the final dataset consisted of 164 patients who had baseline and at least one follow-up time point, and 51 control participants, of whom 43 contributed data at both time points. In total, 642 148 CpG sites passed quality control and were retained for all downstream analyses.

**Supplementary Information S2. Genotype quality control**

Standard quality control procedures were carried out using PLINK (Purcell et al., 2007). Individuals were excluded if they showed >0.02 missingness, heterozygosity rate >|0.20|, or sex mismatch. No individuals met these exclusion criteria. SNPs were removed if they were monomorphic, deviated from Hardy–Weinberg equilibrium (HWE; p < 1×10⁻⁶), showed a missingness >0.02, a difference in missingness between cases and controls >0.02, or a minor allele frequency (MAF) <0.01. Variants with missingness >0.10 during preliminary filtering were also excluded. Autosomal heterozygosity was assessed using linkage-disequilibrium–pruned SNPs (250 kb and 1000 kb windows; r² < 0.1). After all QC steps, 486 010 SNPs remained for downstream analyses.

To control for population stratification, principal component analysis (PCA) was performed using autosomal SNPs after standard quality filters (missingness > 0.01, MAF <0.05, HWE
p < 0.01) and LD pruning (r² < 0.1, 1000 kb window). Individuals deviating more than six standard deviations from the mean on the first five principal components were excluded
(n = 2 patients, n = 3 controls). PCA was then repeated on the cleaned dataset, and the first two principal components were included as covariates in subsequent association analyses.

**Supplementary Information S3. Bacon correction for bias and inflation**

To address potential bias and inflation in our epigenome-wide association study (EWAS), we systematically assessed all models by generating quantile-quantile (QQ) plots and calculating the corresponding inflation factor lambda (λ) based on the original test statistics. The Bayesian correction method implemented in the R package bacon (van Iterson et al., 2017) was used to estimate and correct for bias and inflation in association statistics. Bacon correction was applied exclusively to the model comparing patients and controls, as this was the only model where λ was elevated (λ = 1.55). Input effect sizes and their standard errors were used to fit a hierarchical mixture model estimating parameters for bias and inflation in the summary statistics. The resulting estimates were used to adjust Z-scores, which were subsequently back-transformed into corrected effect sizes and p-values. This procedure addresses systematic confounding and heterogeneity, improving the calibration of test statistics and controlling false positive rates. The effectiveness of the bacon correction was confirmed using QQ plots of the corrected p-values, which demonstrated appropriate calibration (λ = 1.05).

**Supplementary Figures**

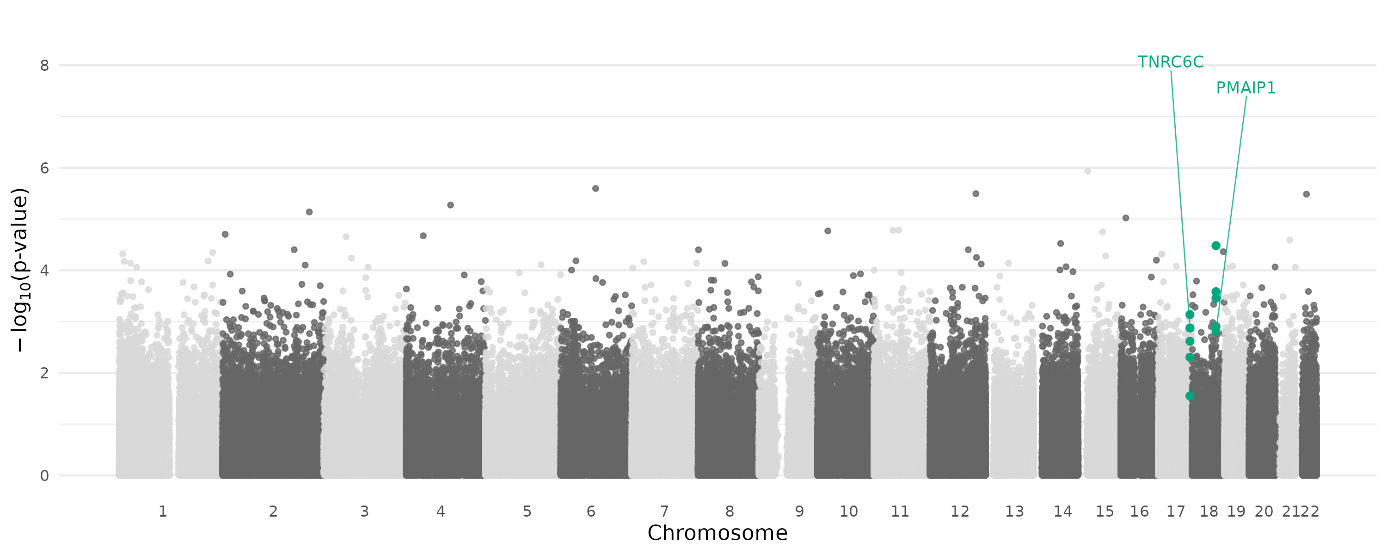

**Supplementary Figure S1: Manhattan plot illustrating the association between DNA methylation and depressive symptom severity at day 28.** Shown are –log₁₀(p-values) across the genome for all tested CpG sites. Differentially methylated regions identified are highlighted in green.

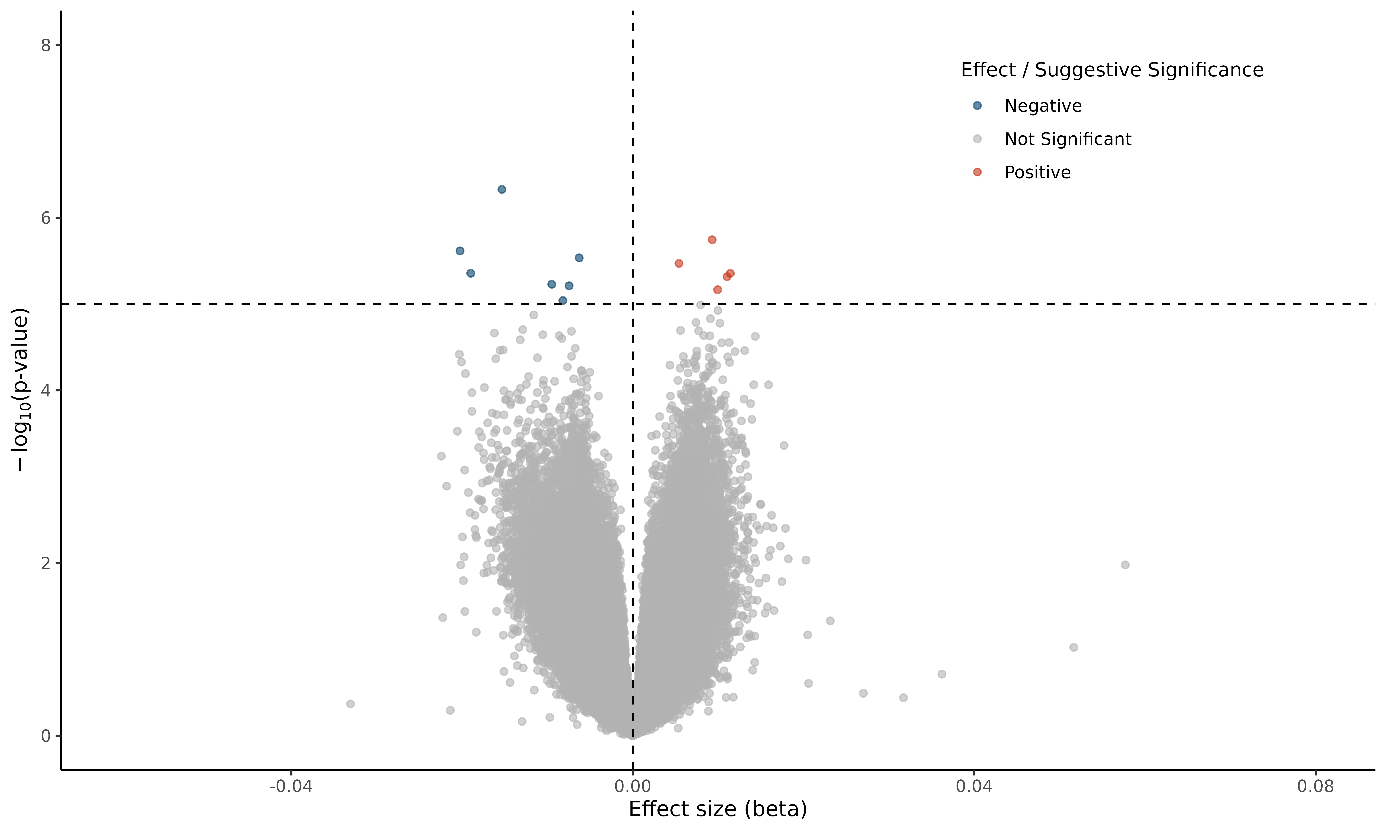

**Supplementary Figure S2. Volcano plot illustrating differential DNA methylation between baseline and day 14 in patients**. Each point represents a CpG site, with the x-axis depicting the estimated change in methylation (β) over time and the y-axis the –log₁₀-transformed p-value. CpG sites showing increased methylation over time are shown in red, and those with decreased methylation are shown in blue. Dashed lines mark thresholds for suggestive significance (p < 1 × 10⁻⁵) and effect size, respectively.

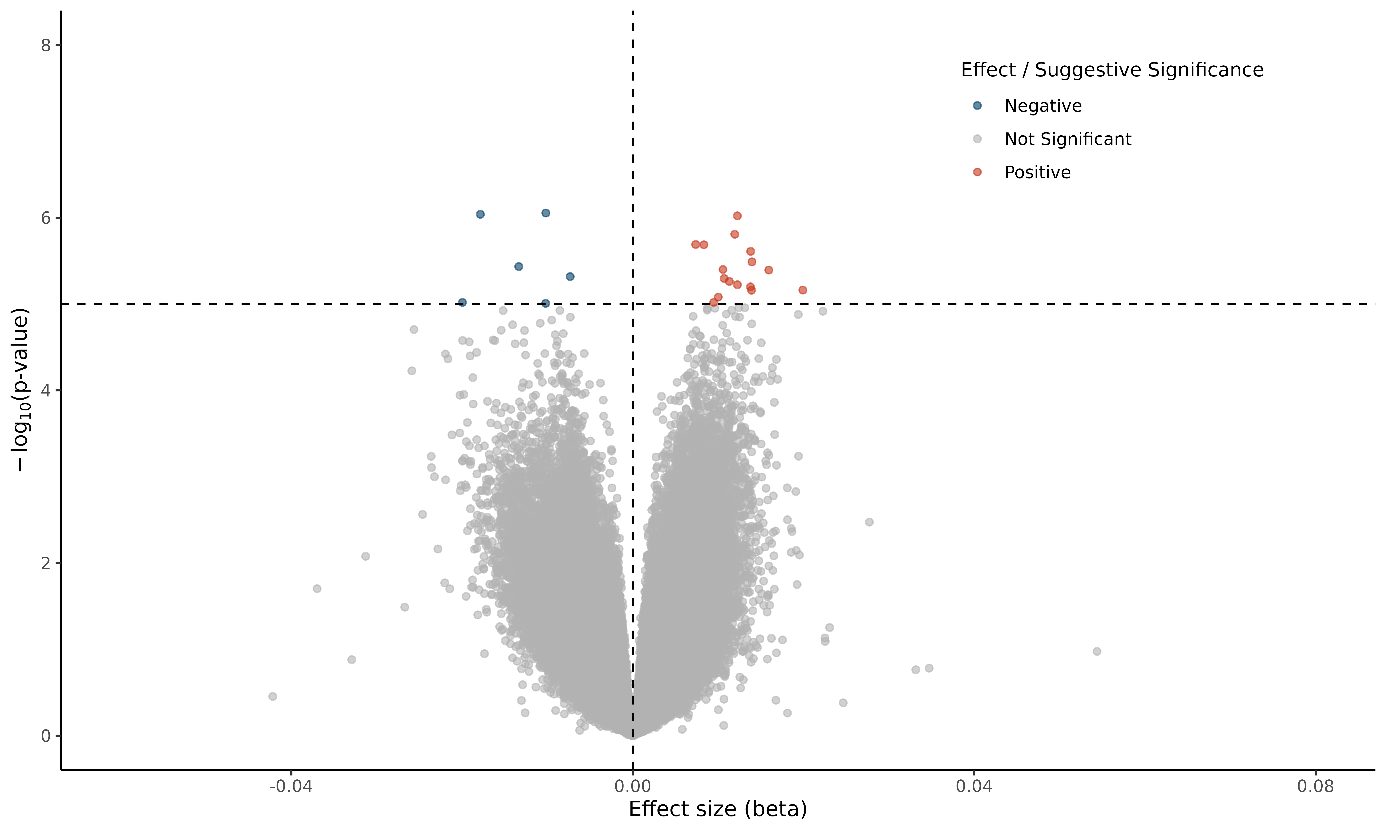

**Supplementary Figure S3. Volcano plot illustrating differential DNA methylation between baseline and day 28 in patients**. Each point represents a CpG site, with the x-axis depicting the estimated change in methylation (β) over time and the y-axis the –log₁₀-transformed p-value. CpG sites showing increased methylation over time are shown in red, and those with decreased methylation are shown in blue. Dashed lines mark thresholds for suggestive significance (p < 1 × 10⁻⁵) and effect size, respectively.

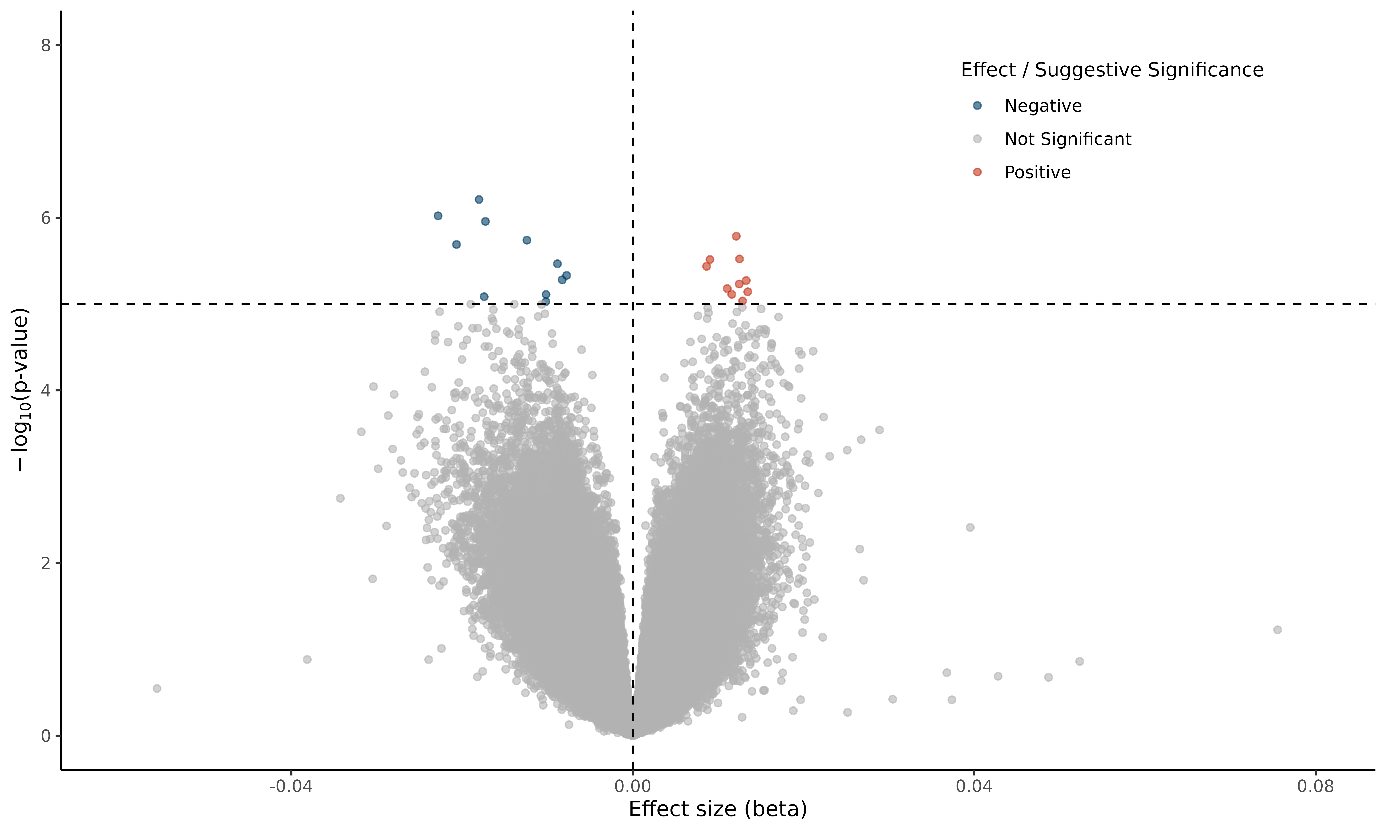

**Supplementary Figure S4. Volcano plot illustrating differential DNA methylation between baseline and day 56 in patients**. Each point represents a CpG site, with the x-axis depicting the estimated change in methylation (β) over time and the y-axis the –log₁₀-transformed p-value. CpG sites showing increased methylation over time are shown in red, and those with decreased methylation are shown in blue. Dashed lines mark thresholds for suggestive significance (p < 1 × 10⁻⁵) and effect size, respectively.
